# Cross-modal integration of reward value during oculomotor planning

**DOI:** 10.1101/613349

**Authors:** Felicia Pei-Hsin Cheng, Adem Saglam, Selina André, Arezoo Pooresmaeili

## Abstract

Reward value guides goal-directed behavior and modulates early sensory processing. Rewarding stimuli are often multisensory but it is not known how reward value is combined across sensory modalities. Here we show that the integration of reward value critically depends on whether the distinct sensory inputs are perceived to emanate from the same multisensory object. We systematically manipulated the congruency in monetary reward values and the relative spatial positions of co-occurring auditory and visual stimuli that served as bimodal distractors during an oculomotor task. The amount of interference induced by the distractors was used as an indicator of their perceptual salience. Our results across two experiments show that when reward value is linked to each modality separately, the value congruence between vision and audition determines the combined salience of the bimodal distractors. However, reward value of vision wins over the value of audition if visual and auditory stimuli have been experienced as belonging to the same audiovisual object prior to the learning of the reward values. The perceived spatial alignment of auditory and visual stimuli is a prerequisite for the integration of their reward values, as no effect of reward value was observed when the two modalities were perceived to be misaligned. These results show that in a task that highly relies on the processing of visual spatial information, the reward values from multiple sensory modalities are integrated with each other, each with their respective weights. This weighting depends on the congruency in reward values, exposure history, and spatial co-localization.

## Introduction

Sensory perception is not merely driven by the incoming sensory inputs but is also affected by top-down information (Gilbert and Sigman, 2007). Among the top-down influences on perception, the effect of reward is particularly important to motivate an agent, facilitate learning, and help the agent to behave adaptively given the limited capacity of both sensory and motor systems. It has been shown that reward acts to modulate selective attention when the subject’s performance was directly linked to the monetary reward (Small et al., 2005; Mohanty et al., 2008; Engelmann et al., 2009), even when the monetary reward was no longer task-relevant (Libera and Chelazzi, 2006; Hickey and van Zoest, 2012; Pooresmaeili et al., 2014; Asutay and Västfjäll, 2016; Luque et al., 2017). Rewards may act as guiding signals for learning and optimizing specific attentional operations (Chelazzi et al., 2013a), and not only can reward increase the salience of associated stimuli (Hickey and van Zoest, 2012), but also enhance the suppression of the distractors (Della Libera and Chelazzi, 2009), change the priority maps of space (Chelazzi et al., 2014), and reduce the intrinsic neural noise in the motor and cognitive control (Manohar et al., 2015).

Nevertheless, although there is a plethora of studies on various aspects of reward, to date, evidence showing the effect of reward-association in one sensory modality on the perception of another modality and, more importantly, how reward from different sensory modalities interact with each other remain scarce. This is despite the fact that in natural environments objects are typically multisensory and comprise multiple attributes that could potentially have either similar or distinct associative values which underscores the importance of understating how information related to reward value is integrated across senses. From a sensory integration point of view, it is well-known that our brain combines multiple sensory signals based on cue integration principles into coherent percepts to reduce uncertainty (Ernst and Banks, 2002; Kersten et al., 2004; Knill and Pouget, 2004), or use information in one sensory modality to prioritize information processing in another sensory modality that better serves as an “expert system” according to the task at hand (Macaluso and Driver, 2005). In other cases, however, perception may be dominated by information from one sensory modality, with other modalities being partially or completely disregarded (Colavita, 1974; Spence, 2009).

Critically, in addition to the physical characteristics of the input stimuli, such as their spatial or temporal properties, recent studies also provided evidence for the effect of cognitive factors on audio-visual integration. For example, expectations of stimulus characteristics can reduce reaction times in a task that requires audio-visual integration (Van Wanrooij et al., 2010a; Zuanazzi and Noppeney, 2018), and emotional (Maiworm et al., 2012) and motivational factors (Bruns et al., 2014) can both influence cross-modal binding processes.

Taken together, the physical and cognitive characteristics of multisensory objects have been shown to affect the processing of information across senses. Although several computational models have been put forward to explain the principles governing cross-modal processing based on the physical characteristics of stimuli (Stein and Meredith, 1993; Calvert, 2001; Stein and Stanford, 2008; Chandrasekaran, 2017), the role of cognitive factors and their possible interactions with physical stimulus characteristics has remained underexplored.

In the present study, our aim was to examine the effect of (1) associated reward values (2) spatial alignment of auditory and visual signals in audiovisual integration. We systematically manipulated these factors during the course of two experiments: Experiment 1 manipulated the congruence of reward values between the auditory and visual stimuli, and Experiment 2 manipulated both reward value congruency and spatial congruency.

The experiments were based on a visually driven saccade task that involved oculomotor interference created by a distractor. Saccades provide a reliable read-out of cross-modal interactions and previous studies have investigated the influence of bimodal targets or distractors on saccade planning (Corneil et al., 2002; Doyle and Walker, 2002; Campbell et al., 2010; Heeman et al., 2016). Importantly, the associated reward value of a visual distractor has been shown to modulate the magnitude of its interference with the planning of the saccades to a target (Hickey and van Zoest, 2012). We hypothesized that in such a task that highly relies on the processing of visual spatial information, visual rewards dominate the effect of bimodal distractors on saccade planning but reward value is integrated across senses if auditory uni-sensory inputs are perceived to have a common source as the visual inputs.

## Material and Methods of Experiment 1

### Participants

Twenty-four participants (18-34 years old; M=25.0, SD=4.0; 11 males) took part in Study 1. One subject was excluded because more than 25% of trials had to be discarded (see the following analyses paragraph for the trial exclusion criteria). All participants were without any neurological or psychiatric disorders and with no recent use of drugs, medications or alcohol dependence, and all had normal or corrected-to-normal sights and normal hearing. Participants gave written informed consent after the experimental procedures were clearly explained to them. They received basic payment plus the payment that was proportional to the accumulated reward value they gained during the reward association part of the experiment. The experiment took approximately 1.5 hours and participants were compensated by €14. The study was conducted in full accordance with the Declaration of Helsinki and was approved by the local Ethics Committee of the Medical University Goettingen (proposal: 15/7/15).

### Apparatus

Eye movements were measured using the Eyelink 1000 eye tracker system (SR Research, Ontario, Canada) in a desktop mount configuration, recording the right eye, with a sampling rate of 1000 Hz. The visual stimuli were presented on a calibrated LCD monitor with a refresh rate of 120 Hz, and the auditory stimuli were presented with over-ear headphones (Sennheiser audiometry headphones, HAD 280). Stimuli were produced by MATLAB and the Psychtoolbox with custom-made scripts.

### Materials and Procedure

Participants were comfortably seated in a dimly lit room, with their head rested on a chinrest. The experiment included a Pavlovian conditioning part to familiarize the participants with the reward associations, and a post-conditioning part that required the participant to perform saccadic eye movements after they had learned the reward associations (**Figure 1A**). Each participant performed 1 block consisting of 120 trials in the conditioning part, 9 training trials to familiarize with the task needed for the post-conditioning part, and 5 blocks consisting of 135 trials (15 repetitions of each of the 9 experimental conditions) for the post-conditioning part. Participants were allowed to take a break for exactly 3 minutes between the blocks. The eye tracker was calibrated at the beginning of the experiment and after each break, by the participant fixating at 13 randomly presented targets spanning the display (EyeLink calibration type HV13).

**Figure 1.**
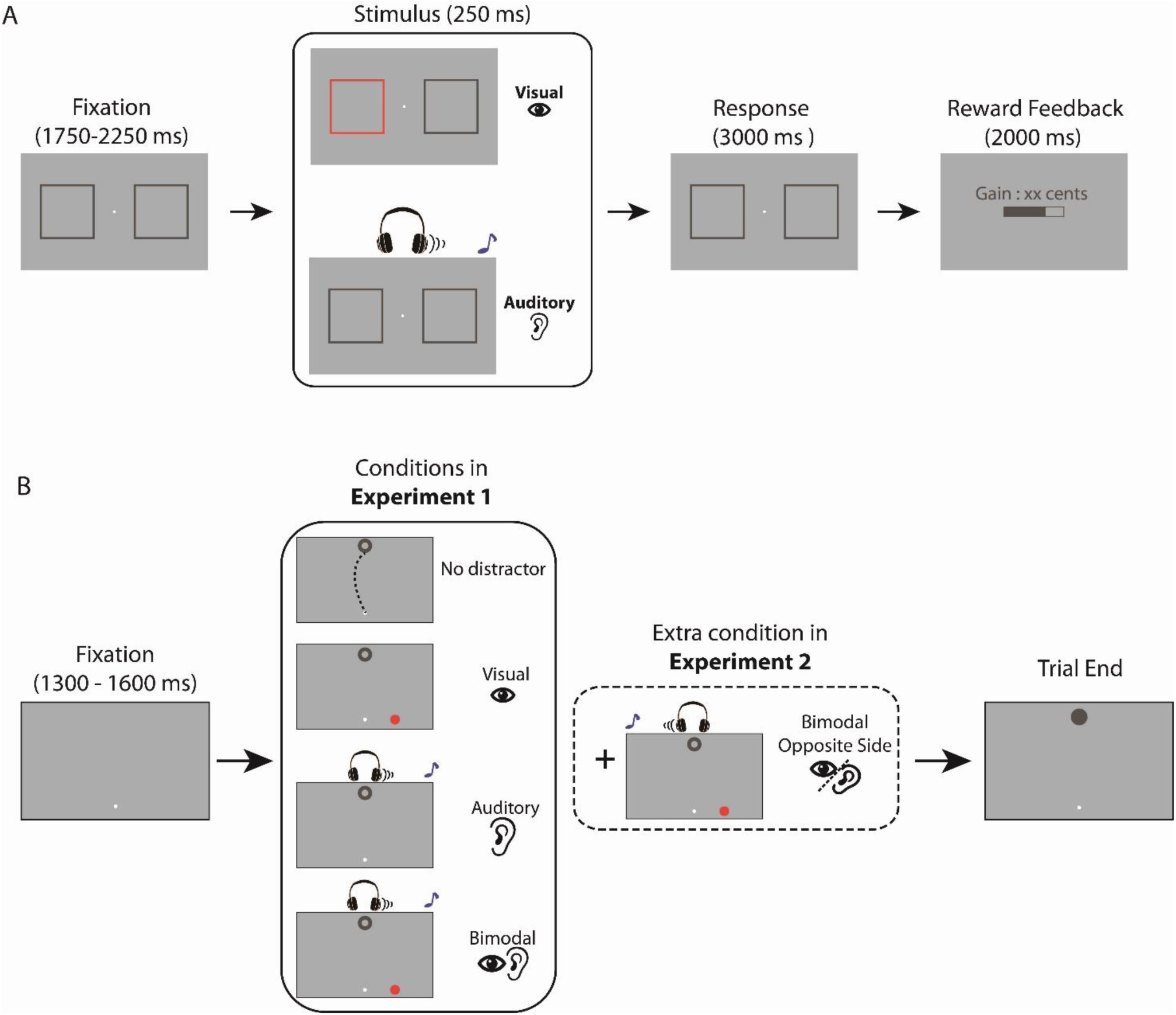
Behavioral tasks. **A. Reward conditioning task**: participants learned the reward associations of 2 colors and 2 sounds through a Pavlovian conditioning paradigm. They were instructed to indicate the side of a sound or a color change by key presses. Correct answers received a non-zero reward, the amount of which depended on the preceding sound pitch (600 or 1000 Hz) or color (magenta or mustard). The reward was displayed graphically as the filled area of a bar as well as a number that corresponded to the amount of monetary reward in Euro cents. Incorrect answers received a reward of zero. **B. Saccadic task in the post-conditioning part of Experiment 1-2**: Participants were instructed to make an eye movement from the fixation point to the target (a ring). In each trial, the participant started by maintaining eye position on the fixation point for a random duration between 1300 and 1600 ms before a target circle was presented either above or below the fixation point, simultaneously with one of the following conditions: (1) with no distractor; (2) with a visual distractor either to the right or left side of the horizontal position of the fixation point; (3) with an auditory distractor presented on the right or left side of the headphone; (4) with a bimodal distractor, which was composed of both the visual and auditory distractors presented on the same side relative to the fixation point. In *Experiment 2*, one more condition was included, in which the auditory distractor was presented on the *opposite side* relative to the visual distractor.

In the conditioning part, participants learned the reward associations of two sounds (600 Hz or 1000 Hz, sawtooth waveform) and two colors (light magenta and light mustard colors, RGB values [171 136 0] and [239 77 255]) by performing a localization task (**Figure 1A**). The loudness level of the sounds and the luminance of the two colors had also been equalized (55 dB and 70 cd/m^2^, respectively), and the background color was mid-grey (RGB: [128 128 128]).

At the beginning of each trial, the participant was instructed to maintain eye position on a dot (0.3°) at the center of the screen for 1750 – 2250 ms. The stimulus display also contained 2 square shaped frames to the left and right side of the fixation point (12° eccentricity, 10° size, RGB: [80 80 80] with a luminance of 70 cd/m^2^). After the fixation period, either a sound (600 Hz or 1000 Hz, counterbalanced across trials) was played on the right or left side of a headphone or a color change occurred in one of the two square frames. All sounds and colors were presented for a fixed duration of 250 ms. Participants were instructed to indicate the correct side of the sound or the color change by pressing the right or left arrow on the keyboard within 3.25 s from the onset of the sounds or colors. Participants received feedback about the amount of obtained reward in each trial, presented graphically as the filled area within a bar as well as a sentence indicating the amount of reward in Euro cents (all with a dark grey color, RGB: [80 80 80]). Incorrect answers led to a reward of zero, whereas correct answers were rewarded by a number drawn from a normal distribution, with high/low mean reward values: 24/4, and a standard deviation of 0.5. The pairing of reward (high or low) with the sound or color was counterbalanced across subjects. Presentation of each modality (sound/color), reward (high/low), and side (right/left side relative to the fixation point) was pseudo-randomized for each subject, and was programmed in a way that none of them repeated consecutively for more than 3 trials. At the end of the conditioning part, the participant was required to indicate the sound and the color that gave higher reward by pressing number ‘1’ or ‘2’ on a keyboard. The orders for the presentation of the 2 sounds and 2 colors were both randomized.

In the post-conditioning part, we used a modified version of the task used by a previous study (Hickey and van Zoest, 2012), where participants had to make an eye movement from the fixation point to a target (**Figure 1B**). The participant first pressed the space bar to begin a trial. Following 1000 ms of successful fixation (eye position maintained within 1° from the center of fixation point with a size of 0.3°), fixation point shrank in size (0.15°) and a second fixation period jittered between 300 and 600 ms had to be fulfilled before the presentation of the experimental stimuli. If eye position offsets more than 1° from the center of fixation point were detected any time within the 1300-1600 ms of the designated fixation period, this interval started over from the beginning. Successful fixations were followed by the presentation of a target (a dark grey ring, RGB: [80 80 80], diameter = 1°) that was presented either 7.38° above or below the fixation point, simultaneously with one of the following conditions: (1) with no distractor; (2) with a visual distractor (colored circle with a radius = 0.8°) either 3.6° to the right or left side of the horizontal position of the fixation point; (3) with an auditory distractor presented on the right or left side of the headphone; (4) with a bimodal distractor, which was composed of both the visual and auditory distractors presented. The visual, auditory and bimodal distractors were based on the stimuli used in the conditioning part, i.e. the colors and sounds that had been associated with reward values. Thus, distractors comprised 9 conditions: no distractor, visual (high or low reward), auditory (high or low reward) and bimodal (high or low visual × high or low auditory, 4 conditions in total). Each condition was repeated 75 times across all trials in a pseudo-randomized sequence. Upon a saccade to the target (within 3° from the center of the target ring), the ring was filled in and stayed in view for another 100 ms before the disappearance of the target and distractors and the start of the next trial.

Same as the conditioning part, after each post-conditioning block, the participant was also required to indicate the sound and the color that gave high reward by pressing the number ‘1’ or ‘2’ on the keyboard and the orders for the presentation of the 2 sounds and 2 colors were both randomized. After the whole experiment, the participants had to fill out a questionnaire regarding the reward values of the stimuli and whether they had any preference towards any stimulus. This questionnaire was used to exclude participants that potentially had misunderstood reward associations (e.g. indicated that high reward sound or color was always presented on the right side).

### Data Analyses

Data were processed and analyzed by using MATLAB (version R2015a). Trials from the training sessions were excluded from the analyses. For each trial, the first saccade after the target onset was analyzed. To detect the saccades, eye position samples in a trial were smoothed using a Savitzky-Golay lowpass filter with an order of 2 and length of 10 ms (Nyström and Holmqvist, 2010). Saccade onsets were defined as the moment when a sample exceeded an angular velocity threshold of 35°/s and an acceleration of 9500°/s^2^. Saccade offsets were calculated as the first sample where the eye position velocity and acceleration fell below the aforementioned thresholds.

Trials with a saccadic latency less than 80 ms or more than 400 ms, with a saccadic duration of more than 120 ms were excluded from the analyses. Additionally, trials where the angular deviation of the saccade endpoint or the saccade amplitude was more than 2.5 standard deviations away from the subjects’ grand mean (across all trials of the session) were defined as outliers and were removed from the analysis. 1 subject was excluded because more than 25% of trials were discarded based on these criteria. In the remaining 23 subjects, 94.78±2.33% of trials were taken into the analysis.

Our statistical inferences and conclusions throughout are based on the mean angular deviation of saccade trajectories, i.e. similar to previous studies of the effect of reward on oculomotor salience (Hickey and van Zoest, 2012). The calculation of the angular deviation of the saccade trajectories for a typical trial is demonstrated in **Figure 2A** and **2B**. Angular deviation was defined as the angular distance between the line joining each sampled eye position along the saccade trajectory to the saccade starting point and the straight path between the saccade starting point and the target. The mean angular deviation is thus the average angular deviation of all samples along the saccade trajectory.

**Figure 2.**
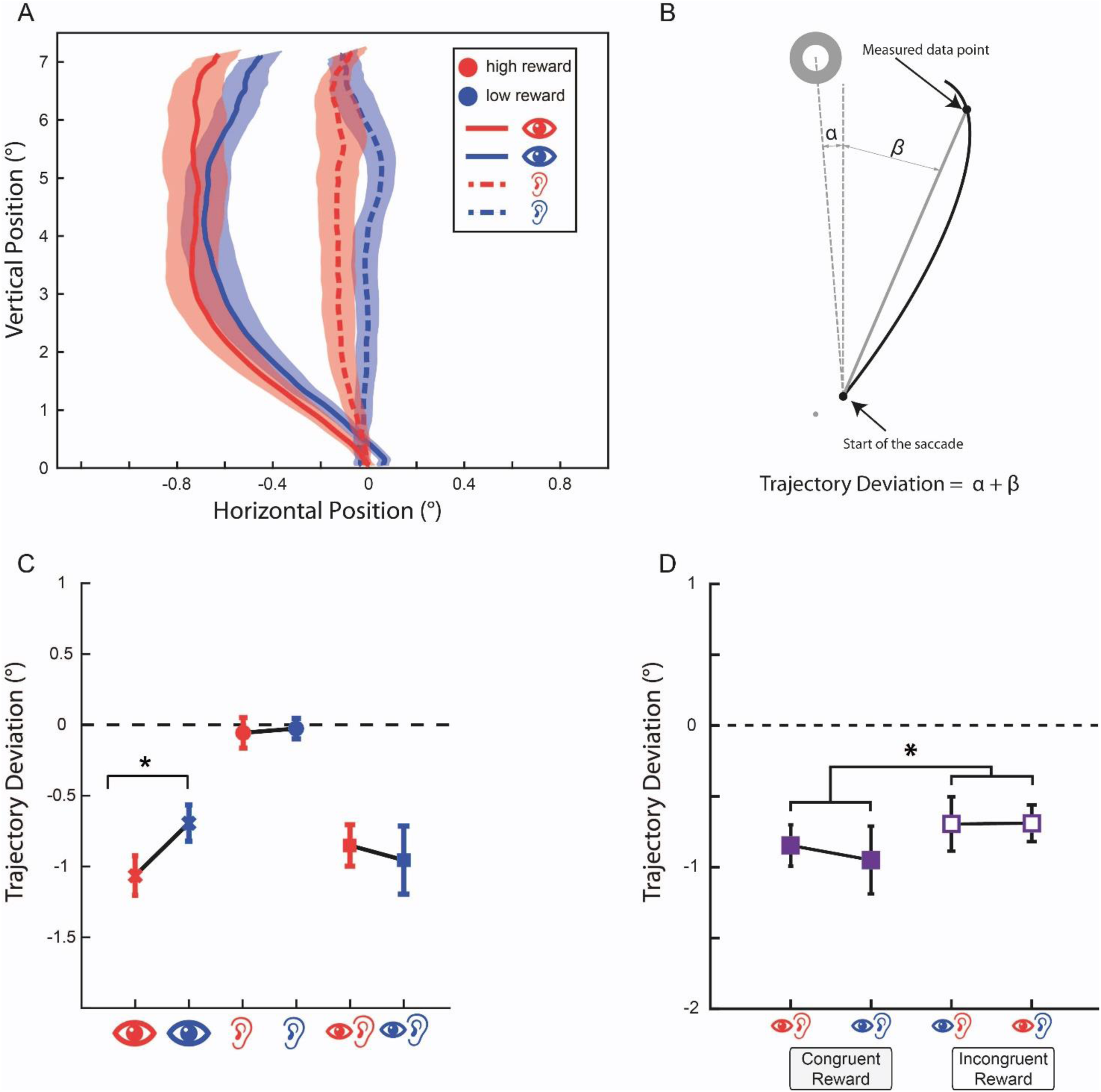
The effect of reward value on saccade trajectories in Experiment 1. **A**. Saccadic trajectories of the high- and low-reward visual and high- and low-reward auditory conditions (indicated by corresponding symbols and colors). **B**. Calculation of the angular trajectory deviation of the saccades. **C**. Baseline corrected, average angular deviations of saccade trajectories as shown in A and B served as a measure of oculomotor interference created by the visual, auditory and bimodal distractors with either high or low reward value (only bimodal distractors with congruent reward values are shown in **C**). Positive values indicate that the saccadic trajectories deviate towards the distractor while negative values indicate deviations away from the distractor. Visual distractors associated with the high reward value resulted in significantly larger deviations away from the distractor, compared to low value distractors. **D**. Baseline corrected saccadic trajectory deviations in all bimodal conditions. From left to right: congruent-reward/both high-value, congruent-reward/both low-value, incongruent-reward/visual low-value & auditory high-value, and incongruent-reward/ visual high-value & auditory low-value conditions. Saccade trajectories had significantly larger deviations when reward values were congruent compared to when they were incongruent across modalities. Error bars indicate the standard error of the mean (s.e.m).

Since the target position could be either in the upper or lower hemi-field, and the distractor could be either on the left or right side relative to the fixation point, in order to examine the angular deviation, we rectified the target position to the upper hemi-field, and the distractor to the right side relative to the fixation point. In each participant, the no-distractor condition served as baseline, and their averaged angular deviation was subtracted from the average angular deviation of each distractor condition with the corresponding target location (i.e. in upper or lower hemi-field). Even in the absence of distractors, saccades may exhibit trajectory deviations which are predominantly towards one of the quadrants. Subtraction of the no-distractor trajectories remedies this directional bias. However, our estimation of the baseline using no-distractor condition could be noisy. Therefore, the baseline was determined by averaging the angular deviation of no-distractor trials which were within 68% confidence interval around the mean in this condition (absolute Z-score <1). We obtained the same statistical results as reported here, when the data was analyzed without the subtraction of the baseline.

We analyzed the saccade average angular deviations by carrying out a series of repeated measures analysis of variance (RANOVA). The first analysis focused merely on the physical property of the distractor, regardless of reward association, with factors for *modalities* (visual, auditory, bimodal). Our primary interest lay in the effect of reward, and the reward congruency between the visual and auditory reward. Therefore the second analysis focused on the effect of *reward* (in the cases where reward was unambiguous, i.e. congruent reward for the bimodal conditions), with factors for modalities (visual, auditory, bimodal) and reward (high vs. low); the third analysis focused on *reward congruency* in the bimodal distractors, with factors for visual reward (high vs. low), and auditory reward (high vs. low).

Latencies of the saccades (the time between the onset of the target and distractors and the onset of the first saccade), saccade duration (the interval between the onset and the offset of a saccade) and the distance of the endpoint of the saccades from the target are also reported in Supplementary Tables (S1-S4).

## Results of Experiment 1

### Conditioning Task

In the conditioning part, participants only had to identify the location of the presented color change/sound in each trial. The mean correct responses were 99.6% across 120 trials, with a standard deviation of 0.6% indicating that subjects could accurately localize the colors/sounds. All subjects had learned the reward value associated with each color and sound (based on their responses at the end of conditioning block and the questionnaire data).

### Saccadic Task

Our statistical analyses for saccadic task are based on the mean angular deviation of each condition. Saccade latencies, durations and the distance of the endpoint from the target are reported in Supplementary Tables (S1-S4).

#### 1. Modality

For the analysis of the effect of modality, the data from all reward conditions of each modality (visual, auditory and bimodal) were pooled. The visual distractors were associated with the strongest trajectory deviations (mean ± SD: −0.88±0.56) followed by the bimodal distractors (mean ± SD: −0.79±0.69), and auditory distractors caused the least deviated (mean ± SD: − 0.04±0.36) saccades. A RANOVA with trajectory deviations as the dependent factor and modality as independent factors revealed a main effect of modalities (F(2,44)=29.05, p=9.0×10^-9^). The effect of bimodal distractors was not different from that of visual distractors (paired t-test, t(22)=0.98, p=0.34). Similar results have been observed previously (Heeman et al., 2016), showing that the trajectory deviations caused by bimodal and visual distractors do not differ when the distractor is remote from the target.

#### 2. Reward association in different modalities

For the analysis of the effect of reward in each modality, we only took the conditions where the distractors carried unambiguous reward values (i.e. only reward congruent conditions were included, **Figure 2C**). A 2-way RANOVA with modality and reward as independent factors revealed a main effect of modality (F(2,44)=28.57, p=1.11×10^-8^). The effect of Reward (F(1,22)=0.75, p=0.39) and the interaction between reward and modalities (F(2,44)=2.53, p=0.091) did not reach statistical significance. A further paired-samples t-test on high versus low visual reward showed a significant effect (t(22)=2.63, p=0.0152), with high visual reward condition showing greater deviations compared to low reward (mean ± SD: −1.0638±0.6827 and − 0.6932±0.6273 for high and low visual reward respectively, **Figure 2A** and **2C**), similar to the results observed previously (Hickey and van Zoest, 2012).

#### 3. Reward Congruency

For the analysis of reward congruency, we took all the bimodal conditions and performed a RANOVA with visual reward (High or Low) and auditory reward (High or Low) as independent factors. Note that in this analysis our bimodal conditions are represented as High/High, Low/Low, Low/High and High/Low, with respect to visual or auditory rewards, with congruent conditions represented as High/High or Low/Low (**Figure 1B** and **2D**). The results of RANOVA showed a significant interaction between the visual and auditory rewards (F(1,22)=4.72, p=0.040), indicating a significant effect of reward congruency. As can be seen in **Figure 2D**, saccadic trajectories exhibited stronger deviations away from the distractor when the visual and auditory reward values were congruent (mean ± SD = −0.90±0.79) compared to when they were incongruent (mean ± SD = −0.69±0.66). None of the main effects reached statistical significance (all Fs<1 and all Ps>0.5).

Having shown that learned reward associations of distractors affect the trajectory of visually-guided saccades, in the following Experiment 2 we further examined the interaction between cognitive factors (reward values of the visual and auditory components of the bimodal distractors) and the physical factors (spatial locations of the visual and auditory components) of the distractor in a similar setup. We also included a pre-conditioning part of the experiment to confirm that the effect of reward only exists after learning of the reward associations.

## Material and Methods of Experiment 2

### Participants

The number of the participants was calculated based on the results of Experiment 1 (paired t-test, two-tailed high visual vs. low visual reward values, α=0.05, β=0.8, N required for this power was 32). In total, 36 participants were recruited, 4 participants were excluded (2 participants did not give correct answers to the questions regarding reward association right after the conditioning part, and were replaced by 2 new participants, 1 participant did not complete the second session, 1 participant had poor calibration values). The final sample comprised thirty-two participants (19–37 years old; M=25.3, SD=4.0; 11 males). The experiment was conducted on two consecutive days. On the first day, all participants received €17; on the second day, all participants received basic payment plus the payment that was proportional to the accumulated value they gained during the reward association part of the experiment (with a maximum of €31). All procedures related to the recruitment of the participants were identical to Experiment 1 and complied with the ethical guidelines as described for that experiment.

### Apparatus

Identical to Experiment 1.

### Materials and Procedure

The conditioning task used for the learning of reward associations and the saccadic task of Experiment 2 were identical to that of Experiment 1 (**Figure 1**), but the procedures and experimental conditions were modified as follows. Experiment 2 was conducted on two consecutive days. On the first day of the experiment, participants first performed a control task in which they indicated the location of the stimuli (visual, auditory or bimodal -with sounds on the same or opposite sides relative to the visual stimulus-) by making an eye movement towards their perceived locations. This way we could estimate the perceived location of the target and distractors that were used in the subsequent saccade task. Visual stimuli were presented 7.38° above or below the fixation point (same as the target location in the saccade task) or 3.6° to the right or left side of the fixation point (same as the distractor location in the saccade task) whereas auditory and bimodal stimuli were presented either to the left or to the right side of the fixation point (same as the distractor location in the saccade task). In total, 10 training trials and 200 experimental trials comprising 20 trials per condition and location were collected. The stimuli were identical to those used in the later saccade task (and the same as in Experiment 1) with the exception that the color of the disks were always dark grey (RGB: [80 80 80], same as the target color in Experiment 1 and the sound had a pitch (800 Hz) that was different from those used in the later parts.

The control task was followed by a pre-conditioning part that required the participant to perform the saccadic task prior to the learning of the reward values (**Figure 1B**). Each participant performed 13 training trials to familiarize with the saccade task followed by 10 blocks consisting of 150 trials for the pre-conditioning part (i.e. 1500 trials in total). In the saccadic task, the distractor conditions were one of the following: (1) no distractor; (2) with a visual distractor (0.8°) either 3.6° to the right or left side of the horizontal position of the fixation point; (3) with an auditory distractor presented on the right or left side of the headphone; (4) with a bimodal distractor, which is composed of both the visual and auditory distractors presented on the *same* side; (5) with a bimodal distractor, which is composed of both the visual and auditory distractors presented on the *opposite* side (**Figure 1B**). Note that visual and auditory distractors could have been associated with either high or low reward values. Bimodal distractors comprised 4 conditions with respect to the associated reward value in each modality (i.e. both modalities high, both modalities low, vision high and auditory low and vision low and auditory high reward values). Since bimodal distractors could be on the same or opposite sides, this resulted in a total of 8 bimodal conditions (i.e. 4 reward pairings × 2 sides). Therefore, experimental stimuli comprised 13 conditions, each repeated 120 times (except for the no-distractor condition that was repeated for 60 trials). The number of trials per distractor condition was increased compared to Experiment 1 (120 trials per distractor condition in Experiment 2 compared to 75 trials per condition used in Experiment 1) in order to have a more robust estimation of the saccades’ trajectory deviations.

On the second day of the experiment, participants first performed the reward conditioning task (160 trials, i.e. 40 trials per reward condition in each modality), followed by the saccadic task (post-conditioning part, same procedure and number of trials as the pre-conditioning part). At the end of the session on day 2, participants performed the control task where they indicated the perceived location of the experimental stimuli with eye movements again (10 training trials and 200 trials of the main task identical to day one). This was to test if training on the saccade task alters the perceived location of the target and the distractors (**Figure 3C**).

**Figure 3.**
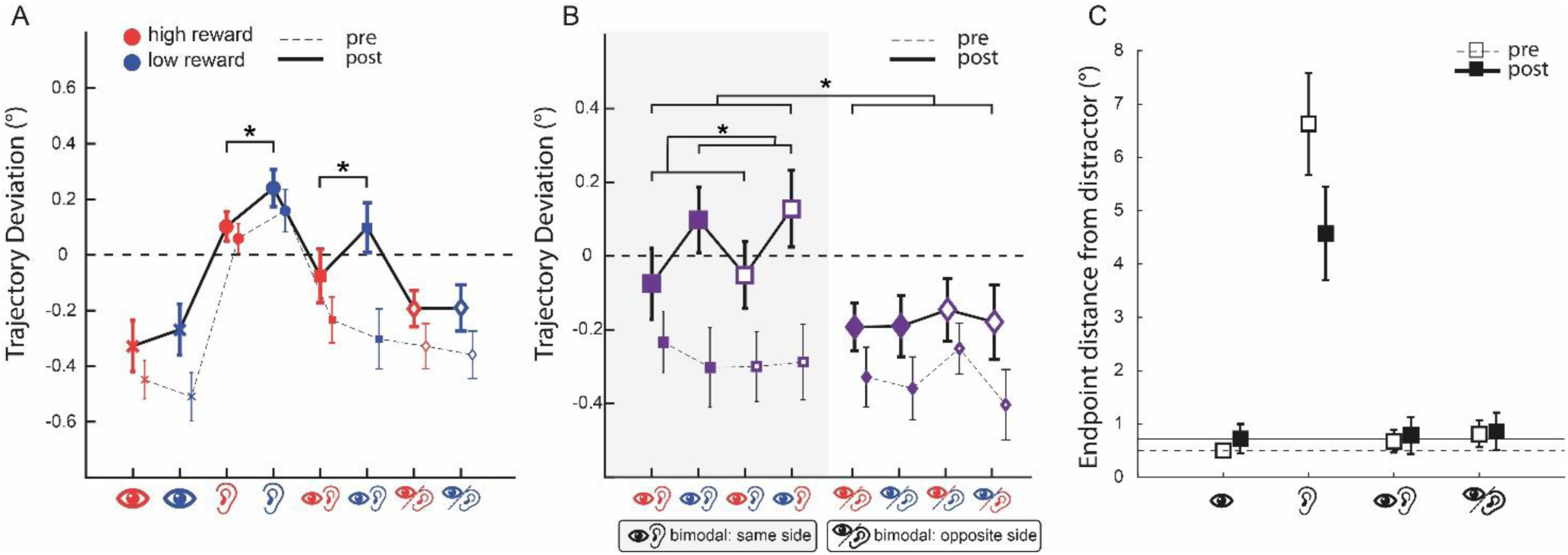
The effect of reward value on saccade trajectories in Experiment 2 and the control task. **A.** Baseline corrected saccadic trajectory deviations in high-versus low-reward value conditions of visual, auditory, bimodal/same-side and bimodal/opposite-sides. The thin error bars and the dashed connecting line between them correspond to the pre-conditioning part, and the thick error bars and the solid connecting line between them indicate the data of the post-conditioning part. The stars indicate significant differences in the post-conditioning part (none of the comparisons reached statistical significance in pre-conditioning part). **B.** Baseline corrected saccadic trajectory deviation in all bimodal conditions. The stars indicate significant effects in the post-conditioning part. Error bars are the standard error of the mean (s.e.m). **C.** The effect of training on the perceived location of auditory and bimodal distractors. The distance of the saccades’ endpoint from the center of visual distractor in control task is displayed for the data collected before participants were exposed to the audiovisual stimuli in the saccade task (white squares) and at the end of the experiment (black squares). Note that in auditory condition no visual distractor was presented and the distance of endpoint is calculated based on the location of distractor across the experiment (i.e. X=±3.6° and Y=0). Dashed and solid horizontal lines are the mean distance of saccades’ endpoints when participants localized visual distractors, before and after exposure to the saccade task respectively. Error bars are the standard error of the mean (s.e.m).

Similar to Experiment 1, we analyzed the mean angular deviations by carrying out a series of RANOVAs. The first analysis focused merely on the physical properties of the distractors, regardless of reward associations, with factors for *modalities* (visual, auditory, bimodal same side, bimodal opposite side); the second analysis focused on the effect of *reward* (in the cases where reward was unambiguous), with factors for modalities (visual, auditory, bimodal same side, bimodal opposite side) and reward (high vs. low); the third analysis focused on contrasting *reward congruency and spatial congruency* in the bimodal distractors, with factors for visual reward (high vs. low), auditory reward (high vs. low), and spatial congruency of the visual and auditory distractors (*same* vs. *opposite*).

### Results of Experiment 2

### Conditioning Part

The mean correct responses were 99.4% across 160 trials, with a standard deviation of 1%.

### Saccadic Task

Based on the exclusion criteria (identical to Experiment 1), in the Pre, 5% of the trials were excluded, and in the Post, 4% of the trials were excluded.

#### 1. The effect of Modality

For the analysis of the effect of modality, high and low visual reward were combined into “Visual”, high and low auditory reward were combined into “Auditory”, all the same-side bimodal conditions were combined into “Same Bimodal”, and all the opposite-side bimodal conditions were combined into “Opposite Bimodal”. Similar to Study 1, a RANOVA revealed a main effect of modalities in both Pre-as well as Post-conditioning parts of the experiment (in Pre, F(3,93)=29.25, p=2.08×10^-13^; in Post, F(3,93)=17.22, p=5.66×10^-9^). Visual distractors caused the most deviated saccadic trajectories away from the distractors (mean ± SD: Pre=-0.47±0.35 and Post=-0.29±0.44), followed by bimodal distractors on the opposite side (Pre=-0.33±0.38 and Post=-0.17±0.33) and bimodal conditions on the same side (Pre=-0.28±0.42 and Post=0.02±0.45). Auditory distractors caused deviations towards the distractors (Pre=0.109±0.309 and Post=0.1714±0.2936). Planned paired t-tests between conditions of interest showed significant differences between saccadic deviations of visual distractors vs. same-side bimodal (Pre: t(31)=3.11, p=0.0040; Post: t(31)=4.85, p=1.45×10^-7^), visual vs. opposite-side bimodal in Pre (Pre: t(31)=2.62, p=0.0135) and in same-side vs. opposite-side bimodal in Post (Post: t(31)=2.37, p=0.0242). Comparison of visual vs. opposite-side bimodal in Post (Post: t(31)=1.69, p=0.10), and same-vs. opposite-side bimodal in pre-conditioning parts (Pre: t(31)=0.94, p=0.35) were not significant. Of particular interest are the results obtained for the comparisons between bimodal conditions and their contrast against visual conditions. Whereas prior to the learning of reward associations no difference between trajectory deviations of bimodal conditions on the same vs. opposite sides was found, after conditioning these conditions diverged from each other, perhaps due to the participants paying more attention to the spatial characteristics of reward-associated sounds and visual stimuli. Bimodal distractors on the same side were associated with significantly smaller deviations compared to visual distractors, which is different from the results obtained in Experiment 1 and indicates that longer training and thus more extensive experience with co-occurring visual and auditory stimuli enhances audiovisual integration, as evidenced by the difference of bimodal condition from both uni-modal conditions. Latencies, durations and the distance of the saccade endpoint from the target are reported in Supplemental Material (Table S2-S4).

#### 2. Reward association in different modalities

In the Pre conditioning part, a RANOVA with modalities and reward as independent factors showed a significant main effect of modality (F(3,93)=25.11, p=5.46×10^-12^) but the effect of reward was not significant (F(1,31)<1 and P>0.7). No significant interaction was found (F(3,93)=1.27, p=0.28).

In the Post conditioning part, however, the RANOVA revealed a main effect of Modalities (F(3,93)=16.06, p=1.72×10^-8^) and a main effect of reward (F(1, 31)=4.26, p=0.047). No interaction was found (F(3,93)=1.07, p=0.36). Therefore, for the Post conditioning part, further paired-samples t-tests were performed: high vs. low auditory reward (mean ± SD: 0.10±0.3 and 0.24±0.37 respectively, t(31)=2.24, p=0.0327), and high vs. low bimodal same-side showed significant differences (mean ± SD: −0.09±0.54 and 0.09±0.50 respectively, t(31)=2.39, p=0.0231), whereas the comparison between high vs. low visual reward (mean ± SD: −0.32±0.51 and −0.26±0.51 respectively, t(31)=-.63, p=0.52) and high vs. low bimodal opposite-side (mean ± SD: −0.19±0.36 and −0.19±0.46 respectively, t(31)=-0.02, p=0.97) was not significant.

#### 3. Spatial congruency and reward congruency

**Figure 3B** shows the results of the *Spatial* versus *Reward Congruency*. A three-way RANOVA with: *Visual Reward* (high vs. low), *Auditory Reward* (high vs. load) and *Spatial Location* (Same vs. Opposite); as factors was performed on the Pre- and Post-conditioning data. In the Pre conditioning part, none of the factors showed a significant difference, and there was no significant interaction either (all Ps>0.1). For the Post-conditioning part, however, the RANOVA showed a significant effect of *Side* (F(1,31)=5.62, p=0.024) and a significant interaction between the *Visual Reward* and *Side* (F(1,31)=5.27, p=0.028). There was a trend for the main effect of visual reward, but this effect did not reach statistical significance (F(1,31)=3.11, p=0.08). All other main and interaction effect had were non-significant (all Fs<1 and Ps>0.5). As can be seen in Figure 3B, the interaction between visual reward and side is driven by a difference between bimodal conditions with high vs. low visual reward that is only present when auditory and visual distractors are on the same side. Accordingly, separate RANOVAs on the trajectory deviations of same-side and opposite-side bimodal distractors showed a significant effect of visual reward only in same-side bimodal distractors (RANOVA, F(1,31), P=0.002), whereas in opposite-side none of the main or interaction effects reached statistical significance (all Fs<1 and Ps>0.5). Therefore, reward value of visual and auditory stimuli was completely disregarded when they were on the opposite sides, whereas the value of visual reward determined the degree of trajectory deviations when the two modalities where perceived to be on the same side.

### Control task: saccadic localization of distractors and the effect of exposure history

The distance between the target and the distractors is an important determinant of the magnitude of distractor interference in the saccadic paradigm that we used in this study. Therefore, in a control task we asked the participants to report the perceived location of the distractors by making a horizontal saccadic eye movement to them. Visual and bimodal distractors were all localized to the same spatial location when participants made saccades to their perceived locations, both before and after experiencing the audiovisual stimuli in the saccade task (mean ± SD of the horizontal position of the saccade endpoints in Pre: 3.57°±0.61°, 3.79°±1.18° and 3.62°±1.45° for visual, bimodal/same-side and bimodal/opposite-side, respectively, and in Post: 3.71°±1.61°, 3.82°±1.98°, 3.77°±2.05°; all Ps>0.1 for paired t-tests). Comparison of the saccades’ accuracy (**Figure 3C**); i.e. the distance of the saccade’s endpoint from the center of the visual distractor revealed a significant difference between bimodal distractors on the same versus different sides at the beginning (mean ± SD in Pre: 0.50°±0.57°, 0.67°±1.19°, 0.81°±1.38°, P= 0.02 in bimodal-same vs. bimodal-opposite, P=0.06 in visual vs. bimodal same, other Ps>0.1). However, this difference ceased to exist after extensive training on the saccade task (mean ± SD in Post: 0.72°±1.57°, 0.78°±1.92°, 0.86°±1.96° for visual, bimodal/same-side and bimodal opposite-side, respectively, all Ps>0.1).

As expected, auditory stimuli were localized less accurately (distance from the center of the visual distractor in Pre: 6.56°±5.34° and in Post: 4.57°±4.87°) and to a location distant from the visual distractor (horizontal position of saccade endpoint in Pre: 9.13°±5.76° and in Post: 6.23°±5.25°). Therefore, across all bimodal and auditory conditions, after extensive training and experience with co-occurring audiovisual stimuli, distractors were localized to a location nearer to the visual distractors. This is similar to the findings of the previous studies (Wozny and Shams, 2011) showing that experience with audiovisual stimuli shifts the localization of auditory signals towards visual stimuli.

### General Discussion (1625 words, 1500 words maximum)

In the present study, our aim was to examine the integration of reward value across sensory modalities during a visually-guided saccade task. More specifically, we examined whether Pavlovian conditioning has an effect similar to the previous studies that utilized transient rewards, and further examine whether factors such as reward congruency and spatial congruency would influence the integration of learned reward values. In Experiment 1, we showed that the reward values learned via Pavlovian conditioning permeated the stimuli even during the non-reward part with a significant effect of reward value on the trajectory deviations of visual distractors and a significant effect of the value congruence in bimodal distractors. In Experiment 2, we re-confirmed the effect of Pavlovian conditioning on the following non-reward part as there was a significant effect of reward value across all modalities. Importantly, after extensive training with bimodal stimuli and experiencing stimuli that were clearly misaligned and those that were not, bimodal distractors with visual and auditory components on the same side were predominantly modulated by the visual reward value. No effect of reward value was observed when visual and auditory distractors were on the opposite sides. The lack of reward-related effects in the pre-conditioning part where no reward association was learned confirms the crucial role of reward learning.

#### Pavlovian conditioning leads to long-lasting changes in perceived salience of reward associated stimuli

Our experimental paradigm was similar to a previous study that specifically examined the impact of transient rewards of visual distractors (i.e. the color of a target associated with high or low reward value in a previous trial could serve as a distractor on next, switch trials) on saccadic trajectories (Hickey and van Zoest, 2012). However, in our study stimulus features were never shared between the target and the distractors, and moreover the distractors acquired their associative values through a separate Pavlovian reward conditioning task. It is therefore, remarkable that completely task-irrelevant distractors that were never rewarded during the oculomotor task could still compete with the processing of the target and lead to changes in the saccade trajectories. Nevertheless, we note that when the number of trials and therefore the duration of the experiment was increased, some of the conditions did not show a significant effect of reward value that is different from previous results (Hickey and van Zoest, 2012) and could be due to a reduction of the reward-related effects over time.

This pattern of result is in line with a series of recent studies showing that associative reward learning can enhance the salience of rewarded stimuli and lead to value-driven attentional capture, even in the following non-reward phase (Theeuwes and Belopolsky, 2012; Yantis et al., 2012; Chelazzi et al., 2013b; Hickey and van Zoest, 2013; Bucker and Theeuwes, 2018; Mine and Saiki, 2018). The impact of associative value during the non-reward phase depends on whether the previously rewarded stimuli serve as targets or distractors. Accordingly, stronger attentional capture by high reward distractors can interfere with the task goals and decrease performance. Our results extend these previous findings to the domain of cross-modal associative reward value.

#### The effect of auditory distractors and their associative reward value on oculomotor planning

Although auditory distractors interfered minimally with the saccade planning compared to visual stimuli, their associative value modulated the responses to co-occurring visual stimuli (in both experiments) and even to uni-sensory auditory stimuli (i.e. significant difference between auditory stimuli with high vs. low reward in Experiment 2). The stronger modulation of saccade trajectories by auditory distractors in Experiment 2 was perhaps due to the longer exposure to the experimental stimuli and recalibration of their perceived positions (Recanzone, 2009; Wozny and Shams, 2011) with a shift to the locations nearer the target, as also demonstrated by the results of our control task (**Figure 3C**).

#### Distinct modes of audiovisual integration: binding by cross-modal reward values versus spatiotemporal alignment

In audiovisual integration, there are several bottom-up and top-down factors that help the brain decide whether audio and visual information should be integrated or segregated (Chen and Spence, 2017). Over decades, numerous studies have shown that stimuli spatial misalignment and temporal asynchrony (Stein and Meredith, 1993) are the main bottom-up factors that disrupt cross-modal integration. Nevertheless, recent studies show emerging evidence that top-down factors such as expectations of stimulus characteristics (Van Wanrooij et al., 2010b), emotional and motivational (Bruns et al., 2014) factors, and semantic congruence (Doehrmann and Naumer, 2008) can all influence audiovisual integration (Macaluso et al., 2016). When incongruent information is provided by auditory and visual signals in the same event, conflicts may be resolved based on their individual modality precision or modality appropriateness (Welch and Warren, 1980), or the system would call for cognitive control (Pessoa, 2009). By juxtaposing the impact of conflicting information related to physical versus cognitive characteristics of auditory and visual stimuli, we could identify two distinct modes of audiovisual integration (**Figure 4**): sensory integration based on the perceived co-localization of auditory and visual inputs (Exp. 2) versus value integration based on the congruency of reward values (Exp. 1) which only takes place if the former mechanism fails to resolve the uncertainty related to the source of uni-sensory signals.

**Figure 4.**
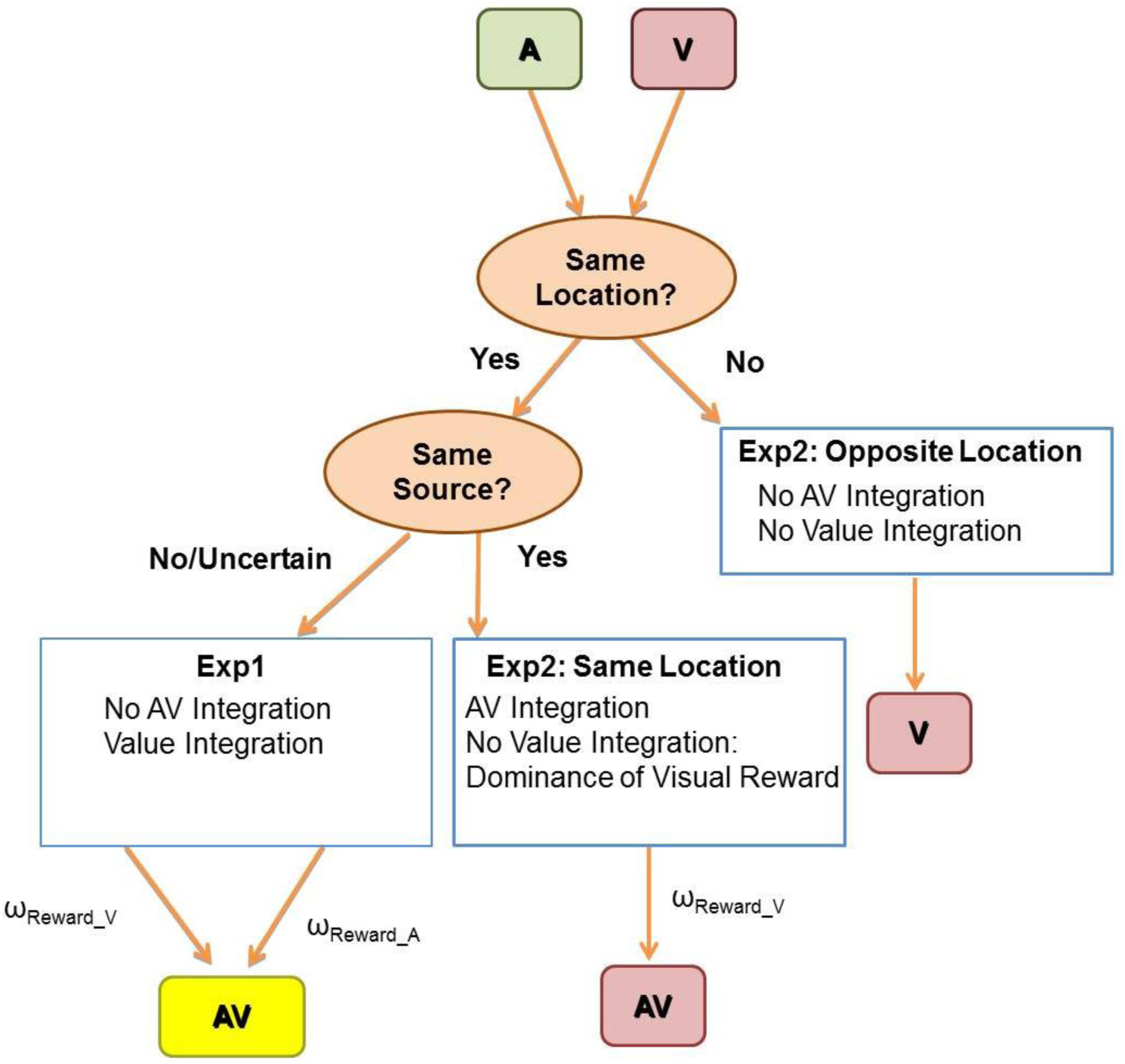
Proposed scheme for how the spatial alignment, reward value and prior experience affect audiovisual integration. In order to decide whether co-occurring auditory (***A***, marked in green) and visual (***V***, marked in red) inputs emanate from the same audiovisual object (***AV***) and should be integrated or not, the first step is to determine whether they share the same spatial location. In case of an obvious misalignment in spatial positions, no further integration occurs, as was the case for bimodal/opposite-sides stimuli in Experiment 2. If ***A*** and ***V*** are assumed to be co-localized, the second step would be to decide whether they have the same source or not. In Experiment 1, the lack of prior exposure to bimodal stimuli led to a greater uncertainty regarding the source of uni-sensory inputs and no audiovisual integration based on sensory features occurred. However, the congruence of reward value was used as an additional information source to weight auditory and visual signals based on the congruency of their reward values (*ω*_*Reward_A*_ and *ω*_*Reward_V*_, for the weight of auditory and visual reward values, respectively). In Experiment 2, prior exposure to audiovisual stimuli and also the experience with overtly misaligned events led to a stronger assumption of relatedness for same-side/bimodal stimuli and their integration at a sensory level (marked as AV integration). However, since in Experiment 2 even apparently misaligned stimuli were sometimes congruent in their reward values, the reliability of value congruence in informing the relatedness of uni-sensory signals was reduced and subsequently more weight was assigned to the reward value in visual modality that also provides more reliable information in a visuospatial task.

Critically, our present study required the participants to learn the associated reward values of the visual and auditory stimuli separately, without them being bound as a unity in the conditioning part. In Experiment 1, participants were asked to perform the saccadic task without prior experience with our bimodal experimental stimuli, therefore the ambiguous spatial location of the auditory stimuli made it difficult to determine whether they are co-localized with the visual stimuli and should be bound into a unity based on their perceived spatial locations. Consequently, in order to determine if visual and auditory stimuli were emanated from the same source, top-down information related to the associated reward values were used to examine the unity assumption, which is similar to cross-modal binding through semantic congruence. This resulted in the significant effect of reward congruency in Experiment 1. Our result is in line with a recent study (Sanz et al., 2018) that also outlined the importance of cross-modal semantic congruency in cross-modal reward integration, and showed that value-driven attentional capture may operate across different sensory modalities for semantically congruent stimuli.

Unlike Experiment 1, in Experiment 2, participants were asked to perform a localization task and the saccadic task *before* the conditioning part; hence they were already exposed to the bimodal stimuli even before learning the reward associations of the stimuli in each modality respectively, and had extensive practice for the saccadic task. Moreover, the contrast between the same-side bimodal and opposite-side bimodal conditions also emphasized the difference between their audiovisual spatial alignments, and could possibly enhance the audiovisual integration for the same-side bimodal conditions. This observation is supported by the fact that in Experiment 1, the saccadic deviation of bimodal conditions did not differ from the visual conditions indicating that auditory stimuli were either localized to a remote location (Heeman et al., 2016) or were assumed to be unrelated to visual stimuli. In Experiment 2, however the saccadic deviation of same-side bimodal conditions significantly differed from the visual and auditory conditions in both pre- and post-parts, indicating the integration of visual and auditory stimuli at a physical level (**Figure 4**). The difference between the perceived spatial alignments of visual and auditory stimuli in Experiment 1 and Experiment 2 is pivotal since in Experiment 1, the auditory and visual stimuli and the information they were associated with could *only* be bound by their reward values, whereas in Experiment 2, it was possible to categorize the spatial locations of the auditory and visual stimuli into the same location in the same-side bimodal conditions, therefore they could be perceived as a single event. Once the auditory and visual stimuli were perceived as a single event, the reward value signaled by the visual stimuli was favored during the saccadic task, as evidenced by the significant interaction between *visual reward* and *side*, due to the fact that the visual system is the “expert system” and contains more reliable information regarding spatial relationships in a visually-driven saccade task. This flexible weighting of reward-related information across sensory modalities is akin to the findings of previous studies that investigated the balance between multisensory integration and uni-sensory dominance by manipulating physical characteristics of stimuli (Yuval-Greenberg and Deouell, 2009).

#### Possible Neural underpinnings

Distractors composed of different sensory modalities had been shown to have an influence on the curvature of the saccades made to a target (Doyle and Walker, 2002; Campbell et al., 2010; Heeman et al., 2016) and we provide evidence for the modulation of these effects by cross-modal reward value. Deviation of saccades towards or away from a visual distractor is proposed to be due to the top-down inhibition of the distractor-evoked responses at the level of Superior Colliculus, a midbrain structure that determines saccade vectors (Munoz et al., 2000). This inhibition could lead to the concurrent inhibition of neural populations that program the saccades to the target and are under a common motor map with the distractor (McPeek et al., 2003; McPeek, 2006). Our current results suggest that cross-modal reward may provide top-down information for such inhibition. Given the dense connectivity between the brain structures that encode reward value in Basal Ganglia and Superior Colliculus (Hikosaka et al., 2014), it is likely that cross-modal integration of reward value occurs at the level of SC. Future studies are needed to explore this possibility.

In conclusion, our results from the two experiments demonstrate that in a saccadic task that highly relies on the processing of visual spatial information, the reward values from a different sensory modality that does not render reliable spatial information can still be integrated with the reward value of the visual modality. The weighting of reward information depends on the assumptions regarding the source of visual and auditory stimuli. Once perceived to be co-localized, the modality that is more task-relevant is assigned with a higher weight for its reward information. In case of uncertainty regarding the spatial co-localization, the congruence of reward value across modalities determines the fate of audiovisual integration.

## Acknowledgements

This work was supported by an ERC Starting Grant (ERC-2016-STG, 716846, *Rewarded Perception*) to AP. We thank Helge Henning and Danilo Postin for their assistance with the data collection.

## Authors’ contributions

FC, AS and AP conceptualized and designed the task. FC, AS and SA conducted the experiments. FC, AS and AP analyzed the data. FC and AP wrote the manuscript. All authors revised the manuscript.

## References

Asutay E, Västfjäll D (2016) Auditory attentional selection is biased by reward cues. Sci Rep 6:36989 Available at: http://dx.doi.org/10.1038/srep36989.

Bruns P, Maiworm M, Röder B (2014) Reward expectation influences audiovisual spatial integration. Attention, Perception, Psychophys 6:1815–1827 Available at: https://doi.org/10.3758/s13414-014-0699-y.

Bucker B, Theeuwes J (2018) Stimulus-driven and goal-driven effects on Pavlovian associative reward learning. Vis cogn 26:131–148 Available at: https://doi.org/10.1080/13506285.2017.1399948.

Calvert GA (2001) Crossmodal Processing in the Human Brain: Insights from Functional Neuroimaging Studies. Cereb Cortex 11:1110–1123 Available at: https://academic.oup.com/cercor/article-lookup/doi/10.1093/cercor/11.12.1110 [Accessed March 31, 2019].

Campbell KL, Al-Aidroos N, Fatt R, Pratt J, Hasher L (2010) The effects of multisensory targets on saccadic trajectory deviations: eliminating age differences. Exp Brain Res 201:385–392 Available at: https://doi.org/10.1007/s00221-009-2045-5.

Chandrasekaran C (2017) Computational principles and models of multisensory integration. Curr Opin Neurobiol 43:25–34 Available at: https://linkinghub.elsevier.com/retrieve/pii/S0959438816302227 [Accessed March 31, 2019].

Chelazzi L, Eštočinová J, Calletti R, Lo Gerfo E, Sani I, Della Libera C, Santandrea E (2014) Altering Spatial Priority Maps via Reward-Based Learning. J Neurosci 34:8594 LP–8604 Available at: http://www.jneurosci.org/content/34/25/8594.abstract.

Chelazzi L, Perlato A, Santandrea E, Della Libera C (2013a) Rewards teach visual selective attention. Vision Res 85:58–72 Available at: https://www.sciencedirect.com/science/article/pii/S0042698912003951 [Accessed August 6, 2018].

Chelazzi L, Perlato A, Santandrea E, Della Libera C (2013b) Rewards teach visual selective attention. Vision Res 85:58–72 Available at: https://www.sciencedirect.com/science/article/pii/S0042698912003951 [Accessed April 2, 2019].

Chen Y-C, Spence C (2017) Assessing the Role of the ‘Unity Assumption’ on Multisensory Integration: A Review. Front Psychol 8:445 Available at: https://www.frontiersin.org/article/10.3389/fpsyg.2017.00445.

Colavita FB (1974) Human sensory dominance. Percept Psychophys 16:409–412 Available at: http://www.springerlink.com/index/10.3758/BF03203962 [Accessed March 31, 2019].

Corneil BD, Van Wanrooij M, Munoz DP, Van Opstal AJ (2002) Auditory-Visual Interactions Subserving Goal-Directed Saccades in a Complex Scene. J Neurophysiol 88:438–454 Available at: https://doi.org/10.1152/jn.2002.88.1.438.

Della Libera C, Chelazzi L (2009) Learning to Attend and to Ignore Is a Matter of Gains and Losses. Psychol Sci 20:778–784 Available at: https://doi.org/10.1111/j.1467-9280.2009.02360.x.

Doehrmann O, Naumer MJ (2008) Semantics and the multisensory brain: How meaning modulates processes of audio-visual integration. Brain Res 1242:136–150.

Doyle MC, Walker R (2002) Multisensory interactions in saccade target selection: Curved saccade trajectories. Exp Brain Res 142:116–130 Available at: https://doi.org/10.1007/s00221-001-0919-2.

Engelmann J, Damaraju E, Padmala S, Pessoa L (2009) Combined effects of attention and motivation on visual task performance: transient and sustained motivational effects. Front Hum Neurosci 3:4 Available at: https://www.frontiersin.org/article/10.3389/neuro.09.004.2009.

Ernst MO, Banks MS (2002) Humans integrate visual and haptic information in a statistically optimal fashion. Nature 415:429 Available at: https://doi.org/10.1038/415429a.

Gilbert CD, Sigman M (2007) Brain States: Top-Down Influences in Sensory Processing. Neuron 54:677–696 Available at: https://www.sciencedirect.com/science/article/pii/S0896627307003765 [Accessed March 20, 2019].

Heeman J, Nijboer TCW, Van der Stoep N, Theeuwes J, Van der Stigchel S (2016) Oculomotor interference of bimodal distractors. Vision Res 123:46–55 Available at: https://www.sciencedirect.com/science/article/pii/S0042698916300207 [Accessed August 3, 2018].

Hickey C, van Zoest W (2012) Reward creates oculomotor salience. Curr Biol 22:R219–R220 Available at: https://www.sciencedirect.com/science/article/pii/S096098221200125X [Accessed July 31, 2018].

Hickey C, van Zoest W (2013) Reward-associated stimuli capture the eyes in spite of strategic attentional set. Vision Res 92:67–74 Available at: http://www.sciencedirect.com/science/article/pii/S0042698913002344.

Hikosaka O, Kim HF, Yasuda M, Yamamoto S (2014) Basal ganglia circuits for reward value-guided behavior. Annu Rev Neurosci 37:289–306 Available at: http://www.ncbi.nlm.nih.gov/pubmed/25032497 [Accessed April 14, 2019].

Kersten D, Mamassian P, Yuille A (2004) Object Perception as Bayesian Inference. Annu Rev Psychol 55:271–304 Available at: https://doi.org/10.1146/annurev.psych.55.090902.142005.

Knill DC, Pouget A (2004) The Bayesian brain: the role of uncertainty in neural coding and computation. Trends Neurosci 27:712–719 Available at: http://www.sciencedirect.com/science/article/pii/S0166223604003352.

Libera C Della, Chelazzi L (2006) Visual Selective Attention and the Effects of Monetary Rewards.

Psychol Sci 17:222–227 Available at: https://doi.org/10.1111/j.1467-9280.2006.01689.x.

Luque D, Beesley T, Morris R, Jack BN, Griffiths O, Whitford T, Le Pelley ME (2017) Goal-directed and habit-like modulations of stimulus processing during reinforcement learning. J Neurosci Available at: http://www.jneurosci.org/content/early/2017/02/13/JNEUROSCI.3205-16.2017.abstract.

Macaluso E, Driver J (2005) Multisensory spatial interactions: a window onto functional integration in the human brain. Trends Neurosci 28:264–271 Available at: http://www.sciencedirect.com/science/article/pii/S0166223605000810.

Macaluso E, Noppeney U, Talsma D, Vercillo T, Hartcher-O’Brien J, Adam R (2016) The Curious Incident of Attention in Multisensory Integration: Bottom-up vs. Top-down. Multisens Res 29:557–583 Available at: https://brill.com/view/journals/msr/29/6-7/article-p557_3.xml [Accessed April 14, 2019].

Maiworm M, Bellantoni M, Spence C, Röder B (2012) When emotional valence modulates audiovisual integration. Attention, Perception, Psychophys 4:1302–1311 Available at: https://doi.org/10.3758/s13414-012-0310-3.

Manohar SG, Chong TT-J, Apps MAJ, Batla A, Stamelou M, Jarman PR, Bhatia KP, Husain M (2015) Reward Pays the Cost of Noise Reduction in Motor and Cognitive Control. Curr Biol 25:1707–1716 Available at: https://www.sciencedirect.com/science/article/pii/S0960982215006120 [Accessed August 6, 2018].

McPeek RM (2006) Incomplete suppression of distractor-related activity in the frontal eye field results in curved saccades. J Neurophysiol 96:2699–2711 Available at: http://www.ncbi.nlm.nih.gov/pubmed/16885521 [Accessed April 14, 2019].

McPeek RM, Han JH, Keller EL (2003) Competition Between Saccade Goals in the Superior Colliculus Produces Saccade Curvature. J Neurophysiol 89:2577–2590 Available at: http://www.physiology.org/doi/10.1152/jn.00657.2002 [Accessed April 14, 2019].

Mine C, Saiki J (2018) Pavlovian reward learning elicits attentional capture by reward-associated stimuli. Attention, Perception, Psychophys 0:1083–1095 Available at: https://doi.org/10.3758/s13414-018-1502-2.

Mohanty A, Gitelman DR, Small DM, Mesulam MM (2008) The Spatial Attention Network Interacts with Limbic and Monoaminergic Systems to Modulate Motivation-Induced Attention Shifts. Cereb Cortex 18:2604–2613 Available at: http://dx.doi.org/10.1093/cercor/bhn021.

Munoz DP, Dorris MC, Paré M, Everling S (2000) On your mark, get set: brainstem circuitry underlying saccadic initiation. Can J Physiol Pharmacol 78:934–944 Available at: http://www.ncbi.nlm.nih.gov/pubmed/11100942 [Accessed April 14, 2019].

Nyström M, Holmqvist K (2010) An adaptive algorithm for fixation, saccade, and glissade detection in eyetracking data. Behav Res Methods 42:188–204 Available at: http://www.springerlink.com/index/10.3758/BRM.42.1.188 [Accessed April 17, 2019].

Pessoa L (2009) How do emotion and motivation direct executive control? Trends Cogn Sci 13:160–166 Available at: http://www.sciencedirect.com/science/article/pii/S1364661309000461.

Pooresmaeili A, FitzGerald THB, Bach DR, Toelch U, Ostendorf F, Dolan RJ (2014) Cross-modal effects of value on perceptual acuity and stimulus encoding. Proc Natl Acad Sci U S A 111:15244–15249 Available at: http://www.ncbi.nlm.nih.gov/pubmed/25288729 [Accessed March 20, 2019].

Recanzone GH (2009) Interactions of auditory and visual stimuli in space and time. Hear Res 258:89–99 Available at: http://www.ncbi.nlm.nih.gov/pubmed/19393306 [Accessed April 14, 2019].

Sanz LRD, Vuilleumier P, Bourgeois A (2018) Cross-modal integration during value-driven attentional capture. Neuropsychologia 120:105–112 Available at: http://www.sciencedirect.com/science/article/pii/S0028393218307231.

Small DM, Gitelman D, Simmons K, Bloise SM, Parrish T, Mesulam M-M (2005) Monetary Incentives Enhance Processing in Brain Regions Mediating Top-down Control of Attention. Cereb Cortex 15:1855–1865 Available at: http://dx.doi.org/10.1093/cercor/bhi063.

Spence C (2009) Explaining the Colavita visual dominance effect. In: Progress in brain research, pp 245–258 Available at: http://www.ncbi.nlm.nih.gov/pubmed/19733761 [Accessed March 31, 2019].

Stein BE, Meredith MA (1993) The merging of the senses. The merging of the senses:xv, 211-xv, 211.

Stein BE, Stanford TR (2008) Multisensory integration: current issues from the perspective of the single neuron. Nat Rev Neurosci 9:255–266 Available at: http://www.ncbi.nlm.nih.gov/pubmed/18354398 [Accessed March 31, 2019].

Theeuwes J, Belopolsky A V (2012) Reward grabs the eye: Oculomotor capture by rewarding stimuli. Vision Res 74:80–85 Available at: http://www.sciencedirect.com/science/article/pii/S0042698912002507.

Van Wanrooij MM, Bremen P, John Van Opstal A (2010a) Acquired prior knowledge modulates audiovisual integration. Eur J Neurosci 31:1763–1771 Available at: https://doi.org/10.1111/j.1460-9568.2010.07198.x.

Van Wanrooij MM, Bremen P, John Van Opstal A (2010b) Acquired prior knowledge modulates audiovisual integration. Eur J Neurosci 31:1763–1771.

Welch RB, Warren DH (1980) Immediate perceptual response to intersensory discrepancy. Psychol Bull 88:638.

Wozny DR, Shams L (2011) Recalibration of auditory space following milliseconds of crossmodal discrepancy. J Neurosci 31:4607 Available at: https://www.ncbi.nlm.nih.gov/pmc/articles/PMC3071751/ [Accessed April 6, 2019].

Yantis S, Anderson BA, Wampler EK, Laurent PA (2012) Reward and attentional control in visual search. Nebr Symp Motiv 59:91–116 Available at: http://www.ncbi.nlm.nih.gov/pubmed/23437631 [Accessed April 11, 2019].

Yuval-Greenberg S, Deouell LY (2009) The dog’s meow: asymmetrical interaction in cross-modal object recognition. Exp Brain Res 193:603–614 Available at: https://doi.org/10.1007/s00221-008-1664-6.

Zuanazzi A, Noppeney U (2018) Additive and interactive effects of spatial attention and expectation on perceptual decisions. Sci Rep 8:6732 Available at: http://www.nature.com/articles/s41598-018-24703-6 [Accessed March 21, 2019].

