## Supplementary Material for "Cross-modal integration of reward value during oculomotor planning"

Table S1. Latencies, durations and the distances of saccades' endpoints from the target in Experiment 1

| | **Visual** | **Auditory** | **Bimodal/Reward Congruent** | **Bimodal/Reward Incongruent (High Reward Visual/Low Reward Auditory)** | **Bimodal/Reward Incongruent (Low Reward Visual/High Reward Auditory)** | **No Distractor** |
|---|---|---|---|---|---|---|
| **Latency (ms) High Reward** | 218.19 ± 23.39 | 188.33 ± 20.98 | 196.94 ± 20.72 | 197.07 ± 19.26 | 198.37 ± 19.58 | 211.27 ± 23.95 |
| **Latency (ms) Low Reward** | 217.41 ± 20.42 | 186.72 ± 20.42 | 201.26 ± 21.82 | | | |
| **Duration (ms) High Reward** | 44.38 ± 33.17 | 44.74 ± 3.37 | 43.91± 3.32 | 43.97 ± 3.24 | 44.63 ± 3.66 | 44.90 ± 3.51 |
| **Duration (ms) Low Reward** | 44.50 ± 3.32 | 44.16 ± 3.33 | 44.26 ± 3.76 | | | |
| **Distance of Endpoint from Target (°) High Reward** | 0.87± .27 | 0.89 ± .32 | 0.91 ± .29 | 0.9 ± 0.29 | 0.91 ± .36 | 0.86 ± 0.24 |
| **Distance of Endpoint from Target (°)Low Reward** | 0.89 ± .28 | .91 ± .29 | 0.90 ± .23 | | | |

Table S2. Latencies of saccades in Experiment 2, before and after learning of reward associations

| | **Visual** | **Auditory** | **Same Side/Congruent Reward/ Bimodal** | **Same Side/ Bimodal (High Reward Visual/Low Reward Auditory)** | **Same Side Bimodal (Low Reward Visual/High Reward Auditory)** | **Opposite Side/Congruent Reward/ Bimodal** | **Opposite Side Bimodal (High Reward Visual/Low Reward Auditory)** | **Same Side Bimodal (Low Reward Visual/High Reward Auditory)** | **No Distractor** |
|---|---|---|---|---|---|---|---|---|---|
| **Latency (ms) High Reward Pre** | 202.08 ± 23.85 | 174.21 ± 19.01 | 182.02 ± 20.24 | 180.63 ± 20.33 | 182.20 ± 20.51 | 178.01 ± 19.93 | 180.60 ± 20.44 | 184.58 ± 21.80 | 193.30 ± 24.16 |
| **Latency (ms) Low Reward Pre** | 204.28 ± 25.95 | 173.35 ± 18.92 | 183.24 ± 20.83 | | | 184.19 ± 21.23 | | | |
| **Latency (ms) High Reward Post** | 180.64 ± 24.28 | 158.97 ± 18.10 | 164.01± 19.38 | 164.92 ± 17.52 | 166.62 ± 19.49 | 164.20 ± 18.73 | 164.68 ± 18.38 | 167.98 ± 19.78 | 193.30 ± 24.16 |
| **Latency (ms) Low Reward Post** | 182.53 ± 24.16 | 159.22 ± 18.2976 | 168.00 ± 20.63 | | | 166.95 ± 19.38 | | | |

Table S3. Durations of saccades in Experiment 2, before and after learning of reward associations

| | **Visual** | **Auditory** | **Same Side/Congruent Reward/ Bimodal** | **Same Side/ Bimodal (High Reward Visual/Low Reward Auditory)** | **Same Side Bimodal (Low Reward Visual/High Reward Auditory)** | **Opposite Side/Congruent Reward/ Bimodal** | **Opposite Side Bimodal (High Reward Visual/Low Reward Auditory)** | **Same Side Bimodal (Low Reward Visual/High Reward Auditory)** | **No Distractor** |
|---|---|---|---|---|---|---|---|---|---|
| **Duration (ms) High Reward Pre** | 43.93 ± 4.00 | 43.73 ± 4.18 | 43.65 ± 4.25 | 43.91 ± 3.99 | 43.68 ± 4.29 | 43.75 ± 4.03 | 43.57 ± 3.93 | 43.88 ± 4.09 | 44.05 ± 4.14 |
| **Duration (ms) Low Reward Pre** | 43.88 ± 4.17 | 43.74 ± 4.30 | 43.60 ± 4.09 | | | 43.92 ± 4.11 | | | |
| **Duration (ms) High Reward Post** | 43.51 ± 3.31 | 43.08 ± 3.76 | 43.32 ± 3.62 | 43.15± 3.38 | 43.35 ± 3.70 | 43.40 ± 3.94 | 43.34 ± 3.44 | 43.35 ± 3.77 | 43.78 ± 3.75 |
| **Duration (ms) Low Reward Post** | 43.66 ± 3.96 | 43.53 ± 4.16 | 43.62 ± 4.06 | | | 43.60 ± 3.84 | | | |

Table S4. The distances of saccades' endpoints from the target in Experiment 2, before and after learning of reward associations

| | **Visual** | **Auditory** | **Same Side/Congruent Reward/ Bimodal** | **Same Side/ Bimodal (High Reward Visual/Low Reward Auditory)** | **Same Side Bimodal (Low Reward Visual/High Reward Auditory)** | **Opposite Side/Congruent Reward/ Bimodal** | **Opposite Side Bimodal (High Reward Visual/Low Reward Auditory)** | **Same Side Bimodal (Low Reward Visual/High Reward Auditory)** | **No Distractor** |
|---|---|---|---|---|---|---|---|---|---|
| **Distance of Endpoint from Target (°) High Reward Pre** | 0.86 ±0.19 | 0.85 ±0.15 | 0.87 ± 0.17 | 0.88 ± 0.19 | 0.84 ± 0.18 | 0.88 ± 0.19 | 0.85 ± 0.16 | 0.89 ± 0.16 | 0.83 ± 0.17 |
| **Distance of Endpoint from Target (°) Low Reward Pre** | 0.85 ± 0.16 | 0.85 ± 0.17 | 0.86 ± 0.17 | | | 0.88 ± 0.19 | | | |
| **Distance of Endpoint from Target (°) High Reward Post** | 0.91 ± 0. 22 | 0.88 ± 0.2 | 0.91 ± 0.22 | 0.88 ± 0.2 | 0.92 ± 0.22 | 0.92 ± 0.21 | 0.89 ± 0.19 | 0.91 ± 0.22 | 0.89 ± 0.20 |
| **Distance of Endpoint from Target (°) Low Reward Post** | 0.91 ± 0. 23 | 0.90 ± .20 | 0.91 ± .21 | | | 0.92 ± .20 | | | |
